# mimicDetector: a pipeline for protein motif mimicry detection in host-pathogen systems

**DOI:** 10.1101/2025.05.02.651971

**Authors:** Kaylee D. Rich, James D. Wasmuth

**Affiliations:** Faculty of Veterinary Medicine, University of Calgary, Calgary, Alberta, Canada; Host-Parasite Interactions Program, University of Calgary, Calgary, Alberta, Canada

**Keywords:** molecular mimicry, host-parasite interactions, sequence search

## Abstract

**Motivation:** Molecular mimicry is a widespread strategy used by pathogens to evade the host immune system and manipulate other host cellular processes. Detecting these events—where pathogen proteins resemble host molecules—is challenging due to limitations in the sensitivity, specificity, and scalability of current bioinformatics tools. The challenges are pronounced when identifying subtle similarities in short protein fragments.

**Results:** We present mimicDetector, an optimized bioinformatic pipeline for systematically identifying protein-level molecular mimicry between pathogens and their hosts. mimicDetector builds on existing *k*-mer-based approaches with three key improvements: (i) improved sensitivity for short-sequence alignments using the PAM30 substitution matrix and tuned BLASTP parameters; (ii) a revised *k*-mer filtering strategy based on bitscore differences rather than percent identity; (iii) the removal of overly conservative homologue exclusion steps. Applied to 17 globally important pathogens, mimicDetector identified a broad and biologically plausible set of mimicry candidates, including helminth proteins mimicking components of the human complement system and a *Leishmania infantum* mimic of Reticulon-4, a regulator of immune cell recruitment.

**Availability and implementation:** mimicDetector is freely available at https://github.com/Kayleerich/mimicDetector/, implemented in Python, and compatible with Unix-based systems.

## 1 Introduction

Many pathogens evade immune detection through molecular mimicry; where pathogen molecules structurally or functionally imitate host molecules. Mimicry has been described across diverse biomolecules, including glycans, lipids, and nucleic acids, with protein mimicry arguably the most widely studied (van Die and Cummings, 2010; Buck *et al*., 2014; Bosch *et al*., 2021; Maguire *et al*., 2024). Identifying mimicry is important not only for understanding how pathogens evade immune surveillance but also for discovering therapeutic targets and revealing co-evolutionary dynamics between hosts and pathogens.

Protein mimicry typically involves short motifs, ranging from five to 20 amino acids. These motifs mediate essential host-pathogen interactions by modulating protein-protein interactions, altering signaling cascades, or directly evading immune detection (as reviewed by Sámano-Sánchez and Gibson (2020)). Despite their biological importance, these short mimic motifs remain challenging to detect computationally. Standard motif discovery tools, such as those from the MEME suite, find overrepresented motifs shared among a predefined set of functionally similar sequences, such as co-regulated genes (Bailey *et al*., 2015).

Unfortunately, when comparing entire proteomes from multiple species without prior functional grouping, these tools are ineffective. Consequently, the hunt for mimic motifs has often relied on local-local sequence alignment tools (Ludin *et al*., 2011; Doxey and McConkey, 2013; Armijos-Jaramillo *et al*., 2021; Emiliani *et al*., 2022). However, these tools are optimised for longer alignments, particularly their default settings, making them poorly suited for capturing subtle similarities in very short alignments (Pearson, 2013).

Previously, we resurrected a *k*-mer-based strategy that aligned pathogen-derived peptides against host and control proteomes and implemented a series of post-alignment filters (Ludin *et al*., 2011; Rich *et al*., 2023). While effective, our approach depended on stringent identity thresholds, potentially missing biologically relevant mimics with subtle sequence divergence. Additionally, homologue removal steps, intended to reduce noise, risked excluding genuine mimics.

Here, we present mimicDetector, an optimised and scalable pipeline for identifying motif mimicry. We systematically evaluate each component of the mimicry detection process— alignment algorithm, scoring matrix, homologue handling, and filtering criteria—to improve sensitivity and specificity in identifying biologically relevant mimic motifs.

## 2 Benchmarking and optimisation

### 2.1 Overall approach

To guide tool selection and parameter optimization for mimicDetector, we designed benchmarking experiments using self-hit and similar-hit peptide datasets derived from a reference pathogen proteome (Figure S1). We selected the proteome of *Plasmodium falciparum* 3D7 (https://www.uniprot.org/proteomes/UP000001450) due to its high annotation quality and representation in previous mimicry studies (Table S1) (Ludin *et al*., 2011; Muthye and Wasmuth, 2023; Yanik *et al*., 2023; Rich *et al*., 2023).

We evaluated the recall performance of five protein alignment search tools: BLASTP, DIAMOND, PHMMER, TOPAZ, and Glam2Scan (Altschul *et al*., 1990; Frith *et al*., 2008; Eddy, 2011; Buchfink *et al*., 2015; Medlar and Holm, 2018). Where possible, we compared two substitution scoring matrices—BLOSUM62 and PAM30—and word sizes—2 and 3—for the seeding step of the alignments. We used query *k*-mers of fixed length (*k*), ranging from 5 to 14 amino acids.

To determine the recall performance of each alignment tool and parameter combination (Table S2), we fragmented each protein in the proteome into overlapping *k*-mers of fixed length (ranging from 5 to 14 amino acids). We built 1000 sets of 1000 sequences randomly sampled from the entire *k*-mer population (Li, 2018). These *k*-mers were aligned back to the original proteome using a given alignment tool. For each *k*-mer, we used a custom python script to determine the number of identical and similar *k*-mers in the original full-length proteins. For the similar *k*-mers, we included up to ‘*k*-3’ mismatches.

### 2.2 Alignment recall

Overall, BLASTP demonstrated the strongest performance across both the identical and similar *k*-mer alignments (Figures S2 and S3). For identical *k*-mer alignments, BLASTP using the PAM30 substitution matrix outperformed BLOSUM62 when *k*≤9 and performed the same when *k*≥10. Reducing the word size from 3 to 2 only improved recall for *k*=5, where seeding is the limiting factor. For similar *k*-mer alignments, BLASTP was again the best performing algorithm, though the choice of scoring matrix was less clearcut and depended on the length of the *k*-mer and the number of mismatches (*m*). PAM30 noticeably outperformed BLOSUM62 when: i) *k*≤9; ii) *k*≥10 and *m*<*k*-9. In contrast, BLOSUM62 slightly outperformed PAM30 for the longer *k*-mers and was more tolerant of fewer mismatches. Reducing the word size provided a slight improvement when using PAM30, but had no discernable effect on the recall of BLOSUM62 alignments.

TOPAZ also performed well for identical *k*-mer alignments. However, it was ultimately removed from consideration for two reasons: a mismatch counting error affected scoring of similar *k*-mer alignments and, even when fixed, the recall of performance of these alignments was lower compared to BLAST, thus limiting its usefulness for detecting biologically plausible motifs. DIAMOND and PHMMER failed to match BLASTP’s performance for identical or similar *k*-mer alignments. Glam2Scan achieved perfect recall for identical *k*-mer alignments but was excluded due to excessive runtimes (Figure S4).

### 2.3 Proteome coverage

In addition to alignment recall, we considered how much of the host proteome was matched by each search strategy with the *P. falciparum k*-mers. We consider it desirable to find a higher coverage of the target proteome, which reduces the likelihood of excluding potentially relevant motifs early in the pipeline. Following the merger of overlapping alignments, we found that BLASTP, PAM30 and wordsize=2 provided the greatest coverage. Further, the coverage flattens when *k*=12 (Figure S5).

### 2.4 Homologue removal

The original pipeline included an early filtering step that removed a pathogen protein that had significant local-local alignments with a host protein, thus excluding its *k*-mers from subsequent searches. One justification for this step—referred to as homologue removal— would be to improve runtimes by reducing the number of *k*-mers searched. However, we considered this approach overly simplistic for two reasons. First, BLASTP is a local alignment tool, so non-homologous regions of the protein could be excluded due to the presence of a promiscuous domain elsewhere on the protein. Second, mimicry can still arise even if the pathogen and host proteins are homologous. For example, the pathogen protein may be one copy of a duplicated gene that has undergone subfunctionalisation and specifically modulates host physiology (Pastrana *et al*., 1998; Armijos-Jaramillo *et al*., 2021;). When we excluded the homologue removal step from the pipeline, there was a small increase in the number of candidate mimic motifs in *P. falciparum* and a marked increased in helminth species (Figure S6).

## 3. Implementation

mimicDetector integrates optimized alignment parameters and refined filtering strategies into a streamlined, modular pipeline (Figure 1). We selected the BLASTP search algorithm, with the PAM30 substitution matrix and wordsize=2 (Table S3). The default *k*-mer length is 12 amino acids. A further change to early versions is the use of bitscore over percentage identity to compare *k*-mer alignments.

**Figure 1:**
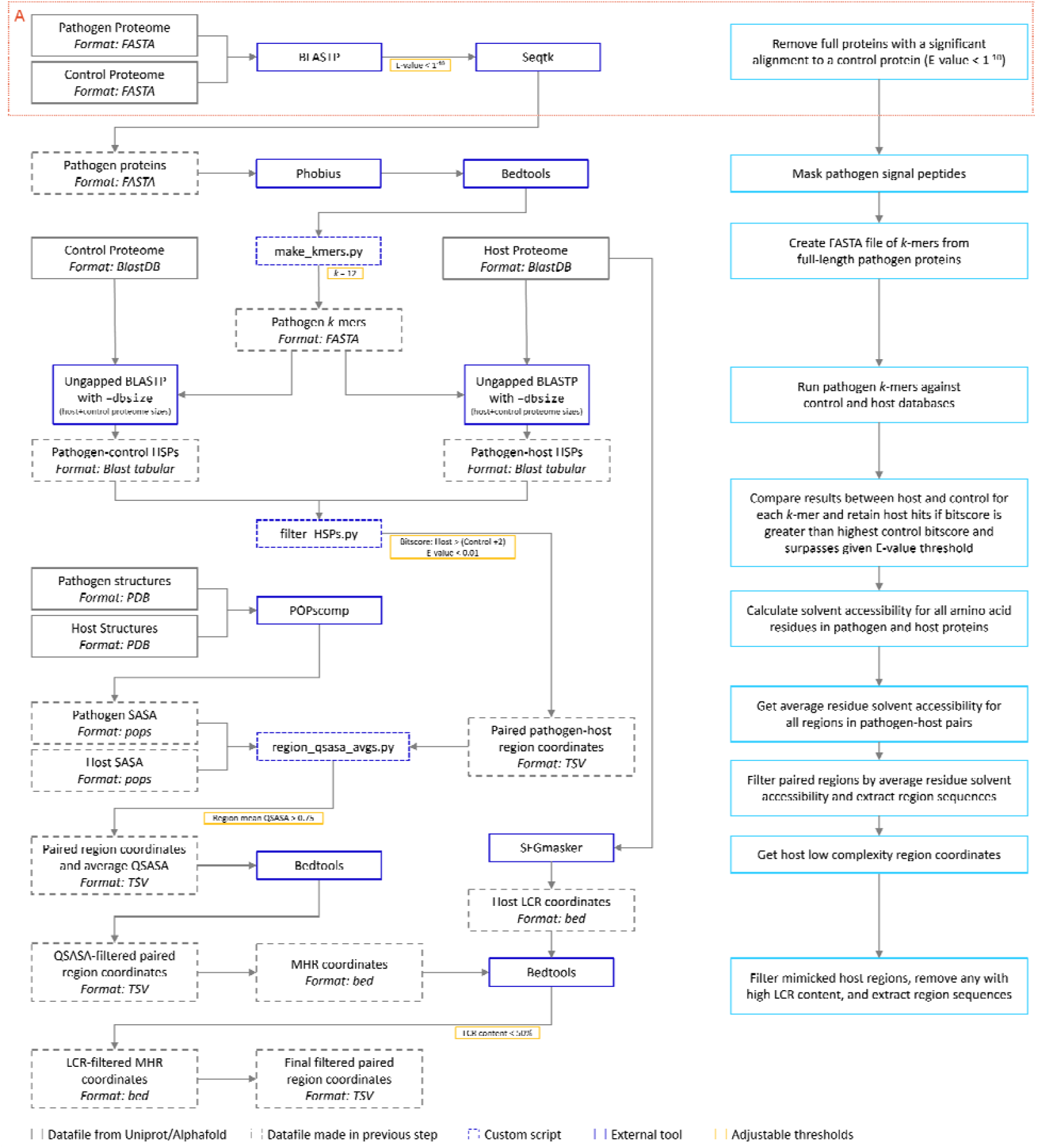
Workflow for mimicDetector pipeline with suggested thresholds. Descriptions for each step are included in the light blue boxes. Box A indicates the homologue removal step, which is not included in the final version of the pipeline

The user begins by supplying proteomes for the pathogen, host, and negative control species (typically non-pathogen) in FASTA format. The pathogen proteome is then fragmented into overlapping *k*-mers, which are used as queries for ungapped local-local alignments.

Post-alignment, mimicDetector applies a sequence of biologically motivated filters. Alignments are retained if the E-value is ≤0.01 and the bitscore of the pathogen-host alignment exceeds that of the best pathogen-control alignment by at least two bits. To reduce false positives, additional filters exclude hits that contain more than 50% low-complexity sequence or less than 75% solvent-accessible residues, determined by Segmasker and POPSCOMP, respectively (Kleinjung and Fraternali, 2005; Wootton and Federhen, 1996).

The final output includes a ranked list of predicted mimicry candidates, annotated with alignment scores, positional information, matched host proteins, and summary statistics. mimicDetector is designed for transparency and flexibility, allowing users to adjust parameters and easily inspect intermediate steps. The pipeline and its accompanying documentation are freely available on GitHub.

## 4. Results and discussion

We applied mimicDetector to a panel of 17 eukaryotic pathogen proteomes to identify potential instances of molecular mimicry targeting the human proteome. A full list of pathogens and control species and their proteome versions is in Table S1. The updated pipeline consistently identified more mimicry candidates than our earlier approach, particularly when homologous proteins were retained and evaluated at the *k*-mer level.

In *Leishmania infantum*, we identified two notable mimics with immunomodulatory potential. One protein, LINF_180015000 (UniProt: A4HXX0), aligned with CD244 (also known as 2B4, UniProt: Q9BZW8), an immune checkpoint receptor expressed on natural killer cells and a target in chronic infection contexts (De Freitas E Silva and Von Stebut, 2021; Sanz *et al*., 2022). A second protein, LINF_310020500 (UniProt: A4I6R4), showed strong alignment to Reticulon-4B (also Nogo-B, UniProt: Q9NQC3), a modulator of leukocyte migration and TLR9 trafficking, which has been implicated in Leishmania-driven changes to macrophage behavior (Kimura *et al*., 2015; Bhattacharya *et al*., 2023).

In parasitic helminths, mimicDetector uncovered mimicry of components of the complement cascade. *Schistosoma mansoni* proteins (UniProt: A0A5K4EZW0, G4V8A8, A0A3Q0KS63 and A0A5K4FDX2) exhibited similarity to C1qA (UniProt: P02745), a subunit of the classical pathway complement component, consistent with known roles for parasite calreticulin in binding and inhibiting C1q function. Similarly, proteins from four species of nematode matched regions of C1qB (UniProt: P02746) (Table S4). These alignments were previously excluded due to homology-based filters, highlighting the improved sensitivity of our new approach. Many helminth species, including those mentioned here, secrete a calreticulin homologue which inhibits classical activation of the complement pathway through interaction with complement protein C1q (Kasper *et al*., 2001; Naresha *et al*., 2009; Floudas *et al*., 2017; Yadav *et al*., 2021; Xian *et al*., 2023; Esperante *et al*., 2023; Jia *et al*., 2024). Functional assays are planned to examine the potential interactions suggested by mimicDetector.

Finally, it is important to note that the search algorithms we tested here were not originally designed for our specific goal. For example, we used Glam2Scan outside of its design parameters and these runtimes are in large part due to formatting the data appropriately for each search. It would be intriguing to explore the use of its underlying Waterman-Eggert algorithm, combined with a scoring matrix, in future mimicry searching (Waterman and Eggert, 1987).

## Supplementary data

See provided supplementary figures

## Data availability

All code and documentation are available at https://github.com/Kayleerich/mimicDetector/. Data related to the results are incorporated into the article and online supplementary material.

