## Supplementary Figures for "mimicDetector: a pipeline for protein motif mimicry detection in host-pathogen systems"

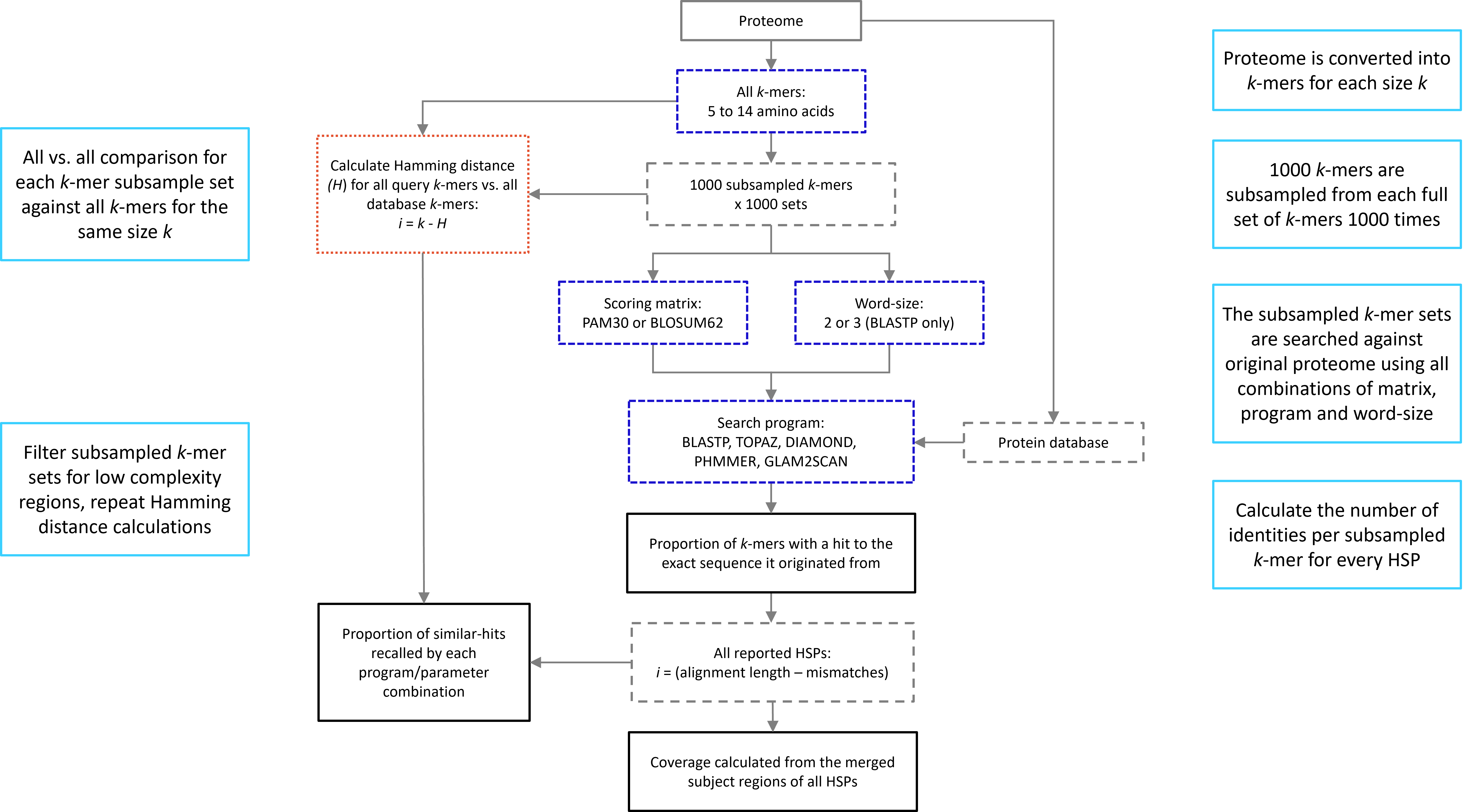

**Figure S1**: Algorithm-parameter sets assessment workflow. Descriptions for each step are included in the light blue boxes. Dark blue boxes indicate the tools or parameters that were assessed. Orange box indicates how true positives were calculated for each subsample.

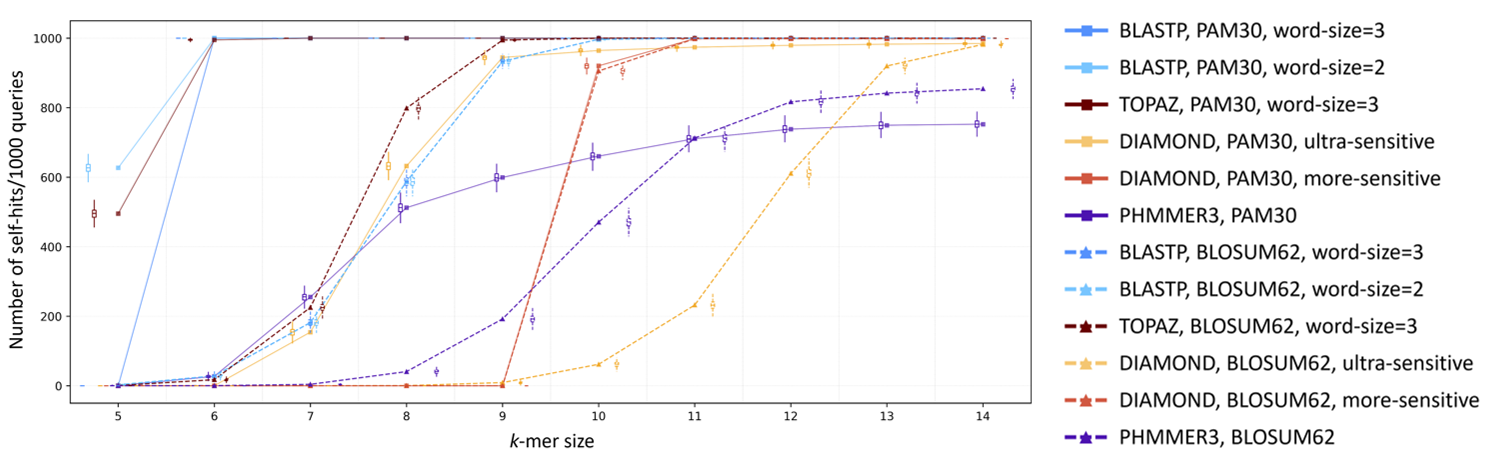

**Figure S2**: Recall of identical hits.

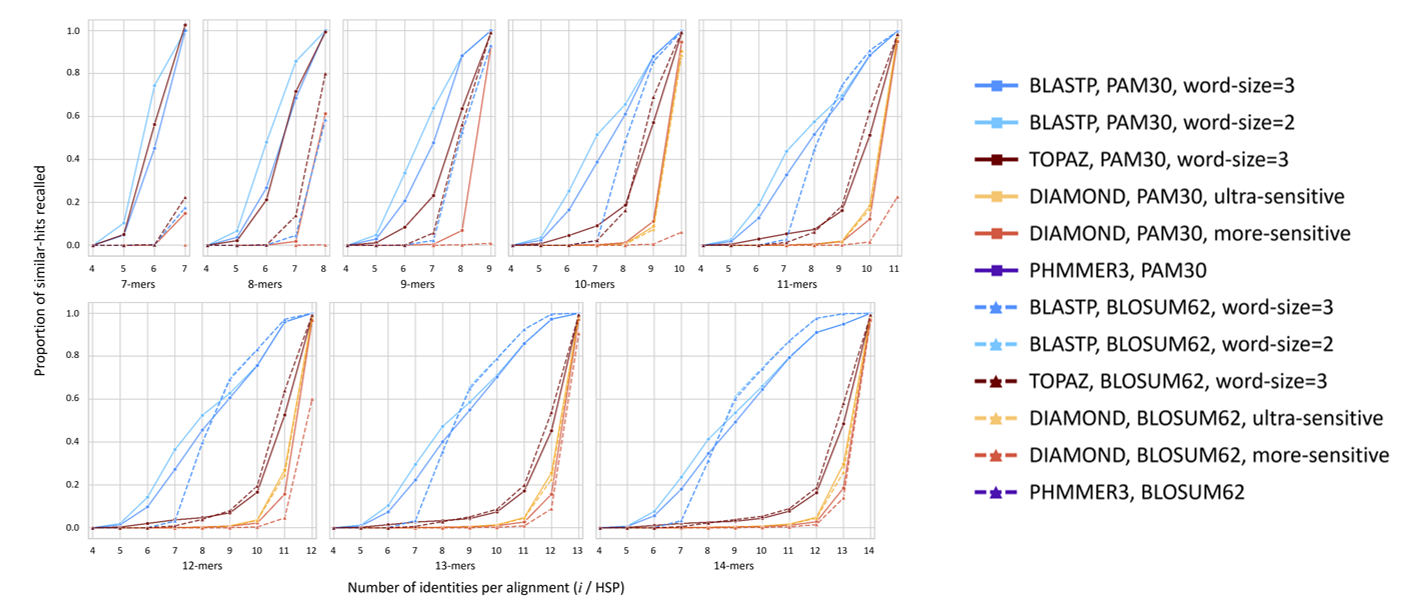

**Figure S3**: Recall of similar hits. Similar-hits were calculated by the proportion of true positives with i identities reported by each tool-parameter combination. True positives were determined through calculation of Hamming distance for each query k-mer. For true positives,

i = k-mer length - Hamming distance; for each tool/parameter combination i = reported alignment length - mismatches for each size k.

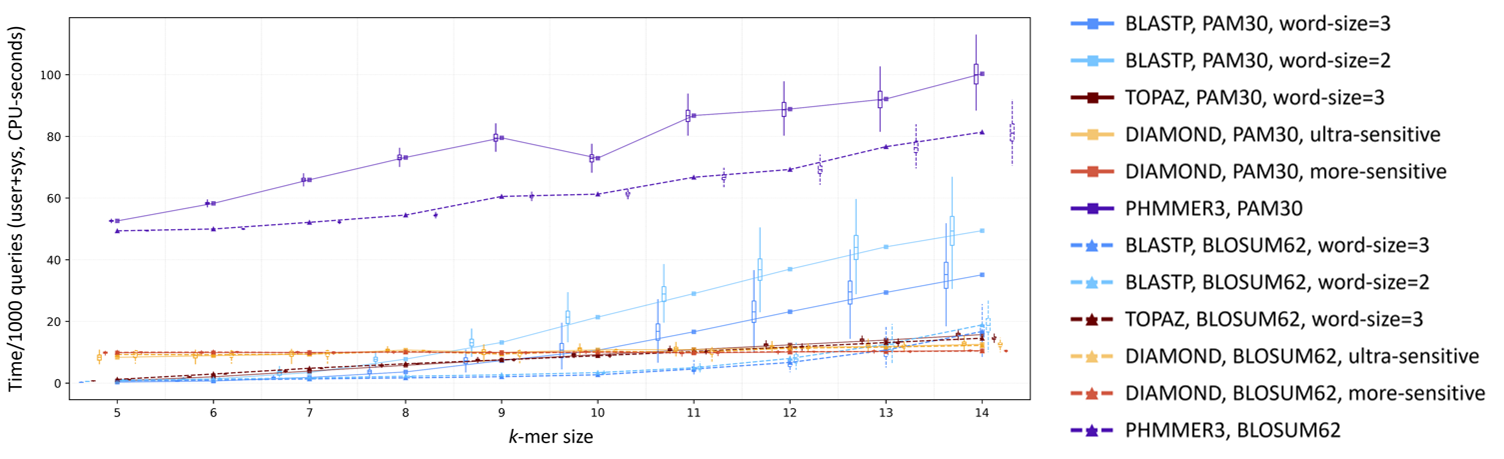

**Figure S4**: Runtime. Time (user + sys CPU time (seconds) used in executing the process) for each tool against the full protein database for each program and k-mer size.

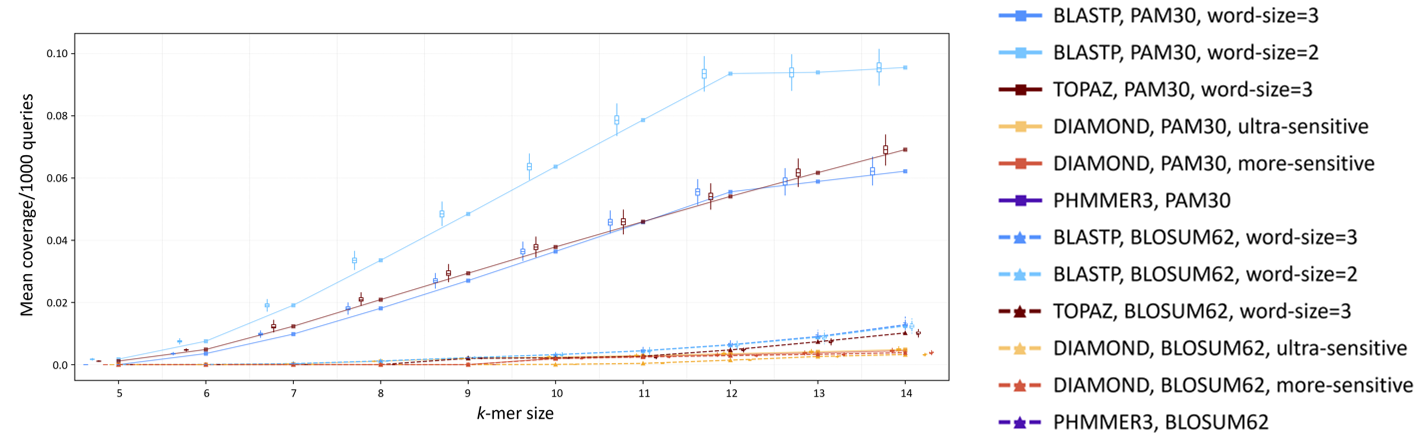

**Figure S5**: Mean coverage of proteome identified by each tool/parameter combination after removing LCRs (query k-mers < 50% content).

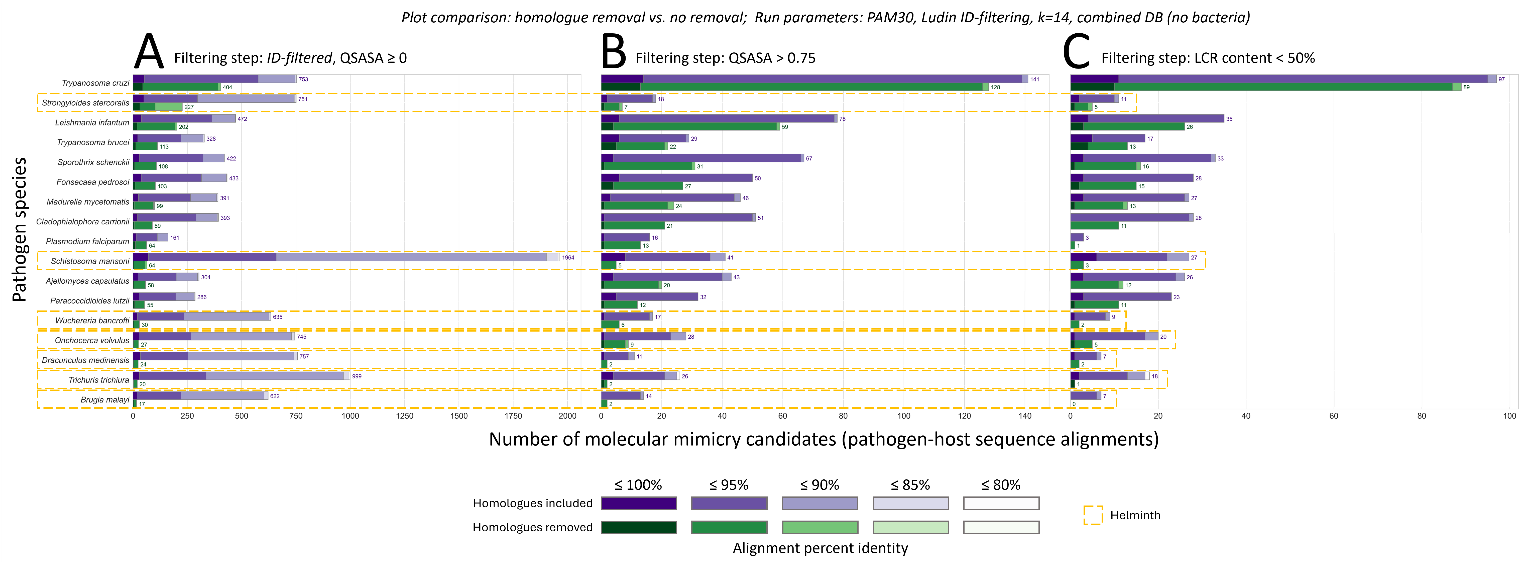

**Figure S6**: Number of mimicry candidates identified for individual species with and without initial homologue filtering. Number of mimicry candidates from runs with homologues included (top bar, purple) and with homologues excluded (bottom bar, green) Helminth species are indicated by dashed boxes to highlight the greater number candidates retained at each filtering stage: A) after BLASTP runs with HSP identity-based filtering, B) after filtering for solvent accessibility, and C) after filtering for low complexity regions.
