## Supplemental Tables for "mimicDetector: a pipeline for protein motif mimicry detection in host-pathogen systems"

**Table S1: Proteomes used in this study.**

| **Species name** | **Species description and relevance** | **Category** | **Uniprot ID** | **# ref. proteins** |
| --- | --- | --- | --- | --- |
| *Homo sapiens* | Human | Host | UP000005640 | 20,577 |
| *Ajellomyces capsulatus* | opportunistic pathogenic yeast, pulmonary histoplasmosis | Pathogen | UP000001631 | 9,214 |
| *Brugia malayi* | filarial nematode, elephantiasis | Pathogen | UP000006672 | 8,825 |
| *Campylobacter jejuni* | gram-negative bacterium, campylobacteriosis | Pathogen | UP000000799 | 1,623 |
| *Cladophialophora carrionii* | melanized fungus, subcutaneous chromoblastomycosis | Pathogen | UP000094526 | 11,173 |
| *Dracunculus medinensis* | guinea worm, dracunculiasis | Pathogen | UP000274756 | 10,868 |
| *Enterococcus faecium* | gram-positive bacterium, opportunistic pathogen with multi-drug antibiotic resistance | Pathogen | UP000325664 | 3,119 |
| *Fonsecaea pedrosoi* | fungus, subcutaneous chromoblastomycosis | Pathogen | UP000053029 | 12,525 |
| *Haemophilus influenzae* | gram-negative bacterium, range of infections including meningitis and pneumonia | Pathogen | UP000000579 | 1,704 |
| *Helicobacter pylori* | gram-negative bacterium, peptic ulcers | Pathogen | UP000000429 | 1,554 |
| *Klebsiella pneumoniae* | gram-negative bacterium, range of healthcare-associated infections | Pathogen | UP000007841 | 5,728 |
| *Leishmania infantum* | protozoan, visceral leishmaniasis | Pathogen | UP000008153 | 8,045 |
| *Madurella mycetomatis* | fungus, mycetoma | Pathogen | UP000078237 | 9,733 |
| *Mycobacterium leprae* | gram-positive bacterium, leprosy | Pathogen | UP000000806 | 1,603 |
| *Mycobacterium tuberculosis* | gram-positive bacterium, tuberculosis | Pathogen | UP000001584 | 3,993 |
| *Mycobacterium ulcerans* | gram-positive bacterium, buruli ulcers | Pathogen | UP000020681 | 9,033 |
| *Neisseria gonorrhoeae* | gram-negative bacterium, gonorrhea | Pathogen | UP000000535 | 2,106 |
| *Nocardia brasiliensis* | gram-positive bacterium, nocardiosis | Pathogen | UP000006304 | 8,414 |
| *Onchocerca volvulus* | filarial nematode, river blindness | Pathogen | UP000024404 | 12,119 |
| *Paracoccidioides lutzii* | fungus, paracoccidioidomycosis | Pathogen | UP000002059 | 8,811 |
| *Plasmodium falciparum* | protozoan, malaria | Pathogen | UP000001450 | 5,376 |
| *Pseudomonas aeruginosa* | gram-negative bacterium, opportunistic pathogen with multi-drug antibiotic resistance | Pathogen | UP000002438 | 5,564 |
| *Salmonella typhimurium* | gram-negative bacterium, range of infections including gastroenteritis | Pathogen | UP000001014 | 4,533 |
| *Schistosoma mansoni* | blood fluke, intestinal schistosomiasis | Pathogen | UP000008854 | 14,097 |
| *Shigella dysenteriae* | gram-negative bacterium, shigellosis | Pathogen | UP000002716 | 3,897 |
| *Sporothrix schenckii* | fungus, sporotrichosis | Pathogen | UP000018087 | 8,673 |
| *Staphylococcus aureus* | gram-positive bacterium, opportunistic pathogen with methicillin resistance | Pathogen | UP000008816 | 2,889 |
| *Streptococcus pneumoniae* | gram-positive bacterium, range of infections including pneumonia | Pathogen | UP000000586 | 2,030 |
| *Strongyloides stercoralis* | threadworm, strongyloidiasis | Pathogen | UP000035681 | 12,823 |
| *Trichuris trichiura* | whipworm, trichuriasis | Pathogen | UP000030665 | 9,625 |
| *Trypanosoma brucei* | protozoan, sleeping sickness | Pathogen | UP000008524 | 8,561 |
| *Trypanosoma cruzi* | protozoan, Chagas disease | Pathogen | UP000002296 | 19,242 |
| *Wuchereria bancrofti* | filarial nematode, lymphatic filariasis | Pathogen | UP000270924 | 13,000 |
| *Arabidopsis thaliana* | thale cress, model organism | Control | UP000006548 | 27,473 |
| *Caenorhabditis elegans* | nematode, model organism | Control | UP000001940 | 19,818 |
| *Candida albicans* | commensal yeast, model organism | Control | UP000000559 | 6,035 |
| *Danio rerio* | zebrafish, model organism | Control | UP000000437 | 25,707 |
| *Dictyostelium discoideum* | slime mold amoeba, model organism | Control | UP000002195 | 12,727 |
| *Drosophila melanogaster* | common fruit fly, model organism | Control | UP000000803 | 13,821 |
| *Escherichia coli* | gram-negative bacterium, model organism | Control | UP000000625 | 4,402 |
| *Glycine max* | soybean, model organism | Control | UP000008827 | 55,855 |
| *Methanocaldococcus jannaschii* | thermophilic methanogenic archaean, model organism | Control | UP000000805 | 1,787 |
| *Oryza sativa* | rice, model organism | Control | UP000059680 | 43,672 |
| *Saccharomyces cerevisiae* | brewer's yeast, model organism | Control | UP000002311 | 6,059 |
| *Schizosaccharomyces pombe* | fission yeast, model organism | Control | UP000002485 | 5,122 |
| *Zea mays* | corn, model organism | Control | UP000007305 | 56,926 |
| *Ciona intestinalis* | sea squirt, model organism | Control | UP000008144 | 16,680 |
| *Trichoplax adhaerens* | placozoan, model organism | Control | UP000009022 | 11,518 |

**Table S2: Parameter sets for optimisation**

| **Algorithm** | **Substitution matrix** | **Word-size** | **Other parameters** |
| --- | --- | --- | --- |
| BLASTP | BLOSUM62 | 2 | -ungapped |
| BLASTP | BLOSUM62 | 3 | -ungapped |
| BLASTP | PAM30 | 2 | -ungapped |
| BLASTP | PAM30 | 3 | -ungapped |
| Diamond BLASTP | BLOSUM62 | NA | --more-sensitive* |
| Diamond BLASTP | BLOSUM62 | NA | --ultra-sensitive* |
| Diamond BLASTP | PAM30 | NA | --more-sensitive* |
| Diamond BLASTP | PAM30 | NA | --ultra-sensitive* |
| PHMMER | BLOSUM62 | NA |  |
| PHMMER | PAM30 | NA |  |
| TOPAZ | BLOSUM62 | 3 | --gapopen 100 |
| TOPAZ | PAM30 | 3 | --gapopen 100 |
| Glam2scan | NA | NA |  |

* gapped alignments removed in post processing

**Table S3: Optimised parameter set**

| **Parameter** | **Value** |
| --- | --- |
| Search algorithm | BLASTP |
| Substitution scoring matrix | PAM30 |
| *k*-mer size | 12 |
| *wordsize* | 2 |
| Scoring thresholds (*b* and *e*)  (pathogen-host vs pathogen-control) | > 2 bits and pathogen-host E-value ≤ 0.01 or E-value ≤ 0.01 if no pathogen-control alignment |
| Solvent accessibility filtering (*q*) | 0.75 |
| Low complexity content (*l*) | 0.50 |

**Table S4: Proteins from parasitic nematodes that share potential mimicry with human C1qB**

| **Species** | **UniprotID** |
| --- | --- |
| *Brugia malayi* | A0A5S6PJ96 |
| *Brugia malayi* | A0A5S6PJ98 |
| *Dracunculus medinensis* | A0A158Q5Z5 |
| *Onchocerca volvulus* | A0A8R1TPF1 |
| *Strongyloides stercoralis* | A0A0K0EJ51 |
